# Modelling locust foraging: How and why food affects hopper band formation

**DOI:** 10.1101/2020.09.21.305896

**Authors:** Fillipe Georgiou, Jerome Buhl, J.E.F. Green, Bishnu Lamichhane, Natalie Thamwattana

**Author notes:** Corresponding author (FG).

## Abstract

Locust swarms are a major threat to agriculture, affecting every continent except Antarctica and impacting the lives of 1 in 10 people. Locusts are short horned grasshoppers that exhibit two behaviour types depending on their local population density. These are; solitarious, where they will actively avoid other locusts, and gregarious where they will seek them out. It is in this gregarious state that locusts can form massive and destructive flying swarms or plagues. However, these swarms are usually preceded by the formation of hopper bands by the juvenile wingless locust nymphs. It is thus important to understand the hopper band formation process to control locust outbreaks.

On longer time-scales, environmental conditions such as rain events synchronize locust lifecycles and can lead to repeated outbreaks. On shorter time-scales, changes in resource distributions at both small and large spatial scales have an effect on locust gregarisation. It is these short time-scale locust-resource relationships and their effect on hopper band formation that are of interest.

In this paper we investigate not only the effect of food on both the formation and characteristics of locust hopper bands but also a possible evolutionary explanation for gregarisation in this context. We do this by deriving a multi-population aggregation equation that includes non-local inter-individual interactions and local inter-individual and food interactions. By performing a series of numerical experiments we find that there exists an optimal food width for locust hopper band formation, and by looking at foraging efficiency within the model framework we uncover a possible evolutionary reason for gregarisation.

**Author summary:** Locusts are short horned grass hoppers that live in two diametrically opposed behavioural states. In the first, solitarious, they will actively avoid other locusts, whereas the second, gregarious, they will actively seek them out. It is in this gregarious state that locusts form the recognisable and destructive flying adult swarms. However, prior to swarm formation juvenile flightless locusts will form marching hopper bands and make their way from food source to food source. Predicting where these hopper bands might form is key to controlling locust outbreaks.

Research has shown that changes in food distributions can affect the transition from solitarious to gregarious. In this paper we construct a mathematical model of locust-locust and locust-food interactions to investigate how and why isolated food distributions affect hopper band formation. Our findings suggest that there is an optimal food width for hopper band formation and that being gregarious increases a locusts ability to forage when food width decreases.

## Introduction

Having plagued mankind for millennia, locust swarms affect every continent except Antarctica and impact the lives of 1 in 10 people [1]. A single locust swarm can contain millions of individuals [2] and is able to move up to 200 kilometres in a day [3]; with each locust being able to consume its own body weight in food [4]. Locusts have played a role in severe famine [5], disease outbreaks [6], and even the toppling of dynasties [7]. More recently, in March 2020 a perfect storm of events caused the worst locust outbreaks in over 25 years in Ethiopia, Somalia and Kenya during the COVID-19 pandemic [8]. Damaging tens of thousands of hectares of croplands and pasture, these outbreaks presented an unprecedented threat to food security and livelihoods in the horn of Africa. In addition, the onset of the rainy season meant the locusts were able to breed in vast numbers raising the possibility of further outbreaks [9].

Locusts are short horned grasshoppers that exhibit density-dependent phase-polyphenism, i.e., two or more distinct phenotype expressions from a single genotype depending on local population density [10]. In locusts there are two key distinct phenotypes, solitarious and gregarious, with the process of transition called gregarisation. Gregarisation affects many aspects of locust morphology from colouration [11], to reproductive features [12], to behaviour [13]. Behaviourally, solitarious locusts are characterised by an active avoidance of other locusts, whereas gregarious locusts are strongly attracted to other locusts. In this gregarious state there is greater predator avoidance on the individual level [14], the group display of aposematic colours has a greater effect of predator deterrence [15], and the resulting aggregations may act as a means of preventing mass disease transmission [16]. It is also in the gregarious state that locusts exhibit large scale and destructive group dynamics with flying swarms of adult locusts being perhaps the most infamous manifestation of this.

Despite the destruction caused by adult swarms, the most crucial phase for locust outbreak detection and control occurs when wingless nymphs form hopper bands, large groups of up to millions of individuals marching in unison [4]. Depending on the species, these groups may adopt frontal or columnar formations, the former being comet like in appearance with dense front and less dense tail [17], and the latter being a network of dense streams [4]. Understanding the group dynamics of gregarious locusts are key to improving locust surveys and control by increasing our ability to understand and predict movement.

In addition to the group dynamics, better knowledge of locust interactions with the environment would help to improve the prediction of outbreaks [18]. On longer time-scales, environmental conditions such as rain events synchronize locust lifecycles and can lead to repeated outbreaks [10]. On shorter time-scales, changes in resource distributions at both small and large spatial scales have an effect on locust gregarisation [19–22]. It is these short time-scale locust-environment interactions that we investigate in this paper, using mathematical modelling to further understand both their effect on swarming and if there is any advantage to gregarisation in this context.

As all the mentioned behaviours arise from simple interactions, understanding the group dynamics of gregarious locusts can also give deep insight into the underlying mechanisms of collective behaviour, consequently they are an important subject of mathematical modelling efforts. Self propelled particles models (SPP) in conjunction with ring shaped arenas [23] and/or field data [24] have been used to extract behavioural characteristics from gregarious locusts [2]. For instance using these techniques Buhl and colleagues have found the critical density for the onset of collective movement [23], the interaction range of locusts (13.5cm), and the way that hopper band directional changes are affected by locust density [17]. One downside of SPP models is that there are few analytical tools available to study their behaviour. In contrast, continuum models can be analysed using an array of tools from the theory of partial differential equations (PDEs). They are most appropriately employed when there are a large number of individuals since they do not account for individual behaviour, instead giving a representation of the average behaviour of the group. The latter (continuum) approach is adopted in this paper.

The non-local aggregation equation, first proposed by Mogilner and Edelstein-Keshet [25], is a common continuum PDE analogue of SPP models [26, 27]. It based a conservative mass system of the form

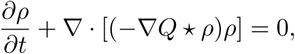

where *Q* is defined as some social interaction potential, *ρ* is the density of species in question, and * is the convolution operation. For this type of model the existence and stability of swarms has been proven [25], and both travelling wave solutions [25] and analytic expressions for the steady states [28] have been found. This model has been further extended to include non-linear local repulsion which leads to compact and bounded solutions [29]. While usually used for single populations, the model has been further adapted to consider multiple interacting species [30].

In a 2012 paper, Topaz et. al. [31] used a multispecies aggregation equation to model locusts as two distinct behavioural sub-populations, solitarious and gregarious. By considering the locust-locust interactions and the transition between the two states, they were able to deduce both the critical density ratio of gregarious locusts that would cause a hopper band to occur and visualised the rapid transition once this density ratio had been reached [31]. For simplicity the model focused on inter-locust interactions and ignored interactions between locusts and the environment. While there exists some continuum models of locust food interactions to investigate the effect of food on peak locust density [32] or to consider hopper band movement [33], we are not aware of any studies that consider locust-locust and locust-food interactions as well as gregarisation in a continuum setting.

The aims of this paper are threefold. Firstly, to derive a mathematical model that is akin the 2012 Topaz model with the inclusion of both locust-food dynamics and local repulsion. Secondly, we use the newly derived model to understand how food interacts with the gregarisation and hopper band formation process. Finally, we investigate under what conditions being gregarious confers an advantage compared to being solitarious.

This paper is organised as follows: we begin with the derivation of a PDE model in which locusts only interact with food when they come into direct contact with it. Our model includes both non-local inter individual interactions and local inter individual and food interactions. Then, we look at some mathematical properties of our model with a homogeneous food distribution. We next use numerical simulations to investigate the effect of food distribution on hopper band formation, and the relative foraging advantage of gregarisation. Finally, we summarise our results and offer ideas for further exploration of the model.

## Models and methods

### Model derivation

In this section we present a PDE model of locust foraging that includes both local inter-individual and food interactions and non-local inter-individual interactions. In this model locusts are represented as a density of individuals (number per unit area) in space and time and are either solitarious, *s*(***x***, *t*), or gregarious *g*(***x***, *t*), with the total local density defined as *ρ*(***x***, *t*) = *s*(***x***, *t*) + *g*(***x***, *t*). For later convenience we will also define the local gregarious mass fraction as

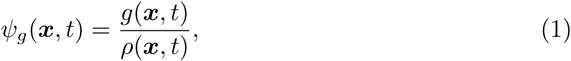

and the global gregarious mass fraction as

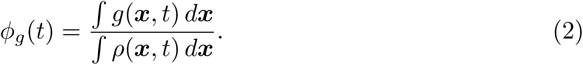

It is possible to write *s* and *g* in terms of Eq (5) as

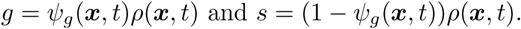

We assume that the time-scale of gregarisation is shorter than the life cycle of locusts, ignoring births and deaths and thus conserving the total number of locusts. We also allow for a transition from solitarious to gregarious and vice-versa depending on the local population density. Hence, conservation laws give equations of the form

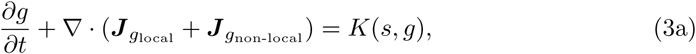

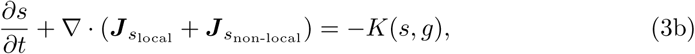

where 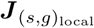 is the flux due to local interactions, 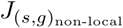 is the flux due to non-local interactions, and *K*(*s, g*) represents the transition between the solitarious and gregarious states.

In addition to locusts we include food resources, let *c*(***x***, *t*) denote the food density (mass of edible material per unit area). We assume that locust food consumption follows the law of mass action and on the time-scale of hopper band formation food production is negligible, giving

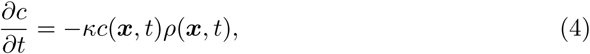

where *κ* is the locust’s food consumption rate.

Finally, for later convenience we will also define the local gregarious mass fraction as

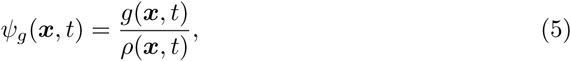

and the global gregarious mass fraction as

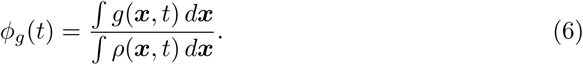

It is possible to write *s* and *g* in terms of Eq (5) as

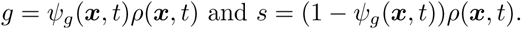

#### Local interactions

We now turn to specifying the local interaction terms in Eq (3a) and Eq (3a). These are captured by taking the continuum limit of a lattice model, we do this by following the work of Painter and Sherratt [34]. We begin by considering solitarious locust movement on a one-dimensional lattice (we assume that local gregarious locust behaviour is the same resulting in a similar derivation). Let 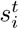 be the number of solitarious locusts at site *i* at time *t*, and let 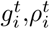, and 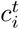 be similarly defined.

We assume that the transition rate for a locust at the *i*^*th*^ site depends on the food density at that site, and the relative population density between the current site and neighbouring sites. If we let 𝒯_*i*_^±^ be the rate at which locusts at site *i* move to the right, +, and left, −, during a timestep, then our transition probabilities are

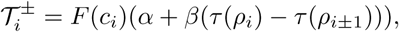

where *F* is a function of food density, *τ* is a function related to the local locust density, and *α* and *β* represent probabilities of movement. If nutrients are abundant at the current site, then we assume locusts are less likely to move to a neighbouring site, which implies *F* is a decreasing function. We set,

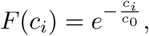

where *c*_0_ is related to how long a locust remains stationary while feeding. We further assume that as the locust population density rises at neighbouring sites relative to the population density of the current site, the probability of moving to those sites decreases proportional to the number of collisions between individuals that would occur. Using the law of mass action, this gives,

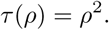

Thus, our transition probabilities are

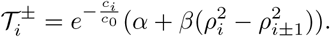

Then the number of individuals in cell *i* at time *t* + Δ*t* is given by

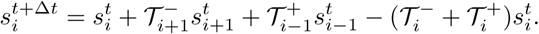

From this, we can derive the continuum limit for both solitarious and gregarious locust densities, and find our local flux as

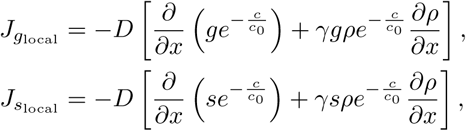

where *D* and *γ* are continuum constants related to *α* and *β* respectively. In higher dimensions, the expressions for fluxes are:

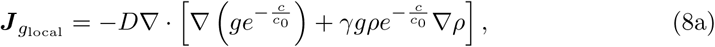

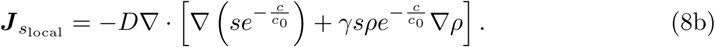

#### Non-local interactions

For our non-local interactions we adopt the fluxes used by Topaz et. al. [31]. By considering each locust subpopulation, solitarious and gregarious, as having different social potentials, we obtain the following expressions for the non-local flux

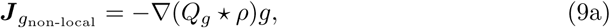

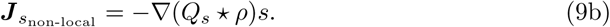

We also adopt the functional forms if the social potentials used by Topaz et. al. [31], as they are used extensively in modelling collective behaviour and are well studied [28]. They are based on the assumption that solitarious locusts have a long range repulsive social potential and gregarious locusts have a long range attractive and a shorter range repulsive social potential. The social potentials are given by,

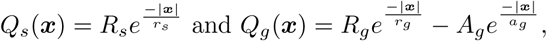

where, *R*_*s*_ and *r*_*s*_ are the solitarious repulsion strength and sensing distance respectively. Similarly, *R*_*g*_ and *r*_*g*_ are the gregarious repulsion strength and sensing distance. Finally *A*_*g*_ and *a*_*g*_ are the gregarious attraction strength and sensing distance.

#### Gregarisation dynamics

For the rates at which locusts become gregarious (or solitarious) we again follow the work of Topaz et. al. [31]. We assume that solitarious locusts transition to gregarious is a function of the local locust density (and vice versa). This gives our equations for kinetics as,

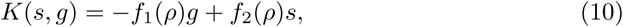

where, *f*_1_(*ρ*) and *f*_2_(*ρ*) are positive functions representing density dependant transition rates. To make our results more directly comparable we again use the same functional forms as Topaz et. al. [31],

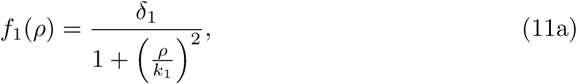

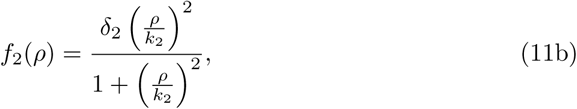

where *δ*_1,2_ are maximal phase transition rates and *k*_1,2_ are the locust densities at which half this maximal transition rate occurs.

#### A system of equations for locust gregarisation including food interactions

By substituting our flux expressions, (8a) through to (9b), and kinetics term (10), into our conservation equations, (3a) and (3b), and rearranging the equation into a advection diffusion system, we obtain the following system of equations

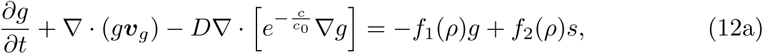

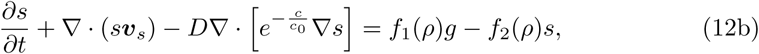

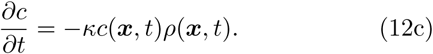

with

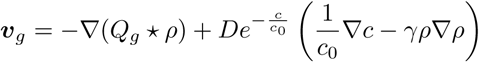

and

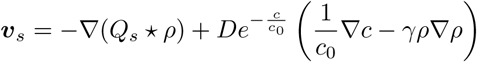

where *f*_1_, *f*_2_, *Q*_*s*_, and *Q*_*g*_ are previously defined.

#### Non-dimensionalisation

We non-dimensionalise Eq (12a), Eq (12b), and Eq (12c), and the explicit expressions for *f*_1_, *f*_2_, *Q*_*s*_, and *Q*_*g*_, using the following scalings

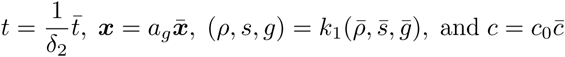

Then dropping the bar notation the dimensionless governing equations are

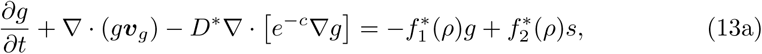

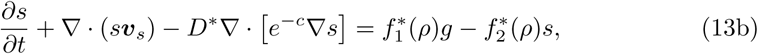

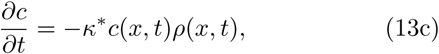

where

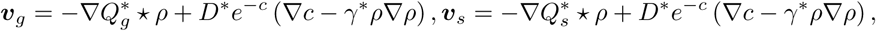

and

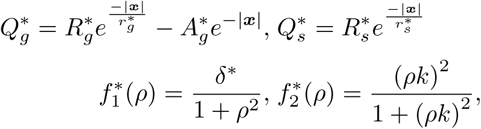

Note that we have introduced the following dimensionless parameters,

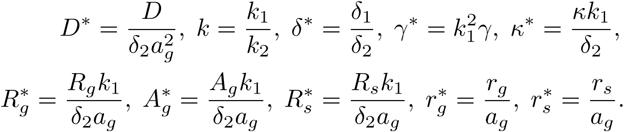

For notational simplicity we drop the ·* notation in the rest of the paper.

## Results

### PDE model analysis

In this section we investigate the behaviour of our model with a spatially uniform food density. Under certain simplifying assumptions, we are able to calculate the maximum density and size of gregarious hopper bands for both small and large numbers of locusts. We then consider the linear stability of the homogeneous steady states to investigate how the availability of food affects hopper band formation, before finally investigating how the center mass is affected by locust interactions. Full details of all calculations can be found in S1 Appendix.

#### Density of gregarious hopper bands

Under some simplifying assumptions we can estimate the maximum density and width of gregarious locusts for both small and large numbers of locusts, termed the small and large mass limits, in one dimension. To begin, we assume that *c* is constant and not depleting, there are minimal solitarious locusts present in the hopper band (i.e. *ρ* ≈ *g*), and the effect of phase transitions in the hopper band is negligible (i.e. *f*_1_(*ρ*)*s* = *f*_2_(*ρ*)*g* = 0). Finally, while the support of *g* is infinite (due to the linear diffusion) the bulk of the mass is contained as a series of aggregations, we will approximate the support of a single aggregation as Ω. Using these assumptions we can rewrite Eq (13a) as a gradient flow of the form,

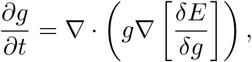

where

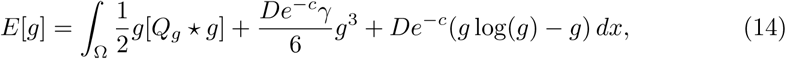

with the minimisers satisfying

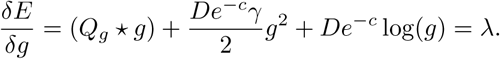

Next, we follow the work of [29, 32, 35] and with a series of simplifying assumptions we consider both the large and small mass limit in turn. First, define the mass of locusts as

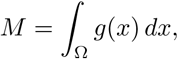

To find the large mass limit, we begin with Eq (14) and assume that *g*(*x*) is approximately rectangular and for a single aggregation that the support is far larger than the range of 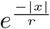. This gives 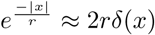 (where *δ*(*x*) is the Dirac delta function), and therefore *Q*_*g*_ ≈ 2 (*R*_*g*_*r*_*g*_ − *A*_*g*_) *δ*(*x*). Using these assumptions we can estimate the maximum gregarious locust density, ||*g*||_*∞*_, as

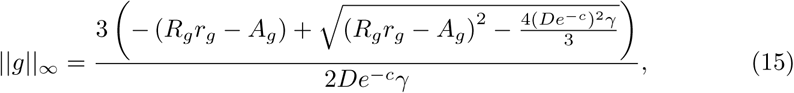

with support,

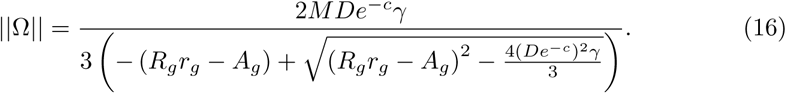

The accuracy of this approximation is illustrated by Fig 1. We observe that within our model as *c* increases so too does the maximum density of our locust formation. However, as the mass of locusts, *M*, increases the maximum density remains constant and the support ||Ω|| becomes larger. Finally, by using these derived relationships with field measurements of maximum locust densities we can estimate values of *γ*.

**Fig 1.**
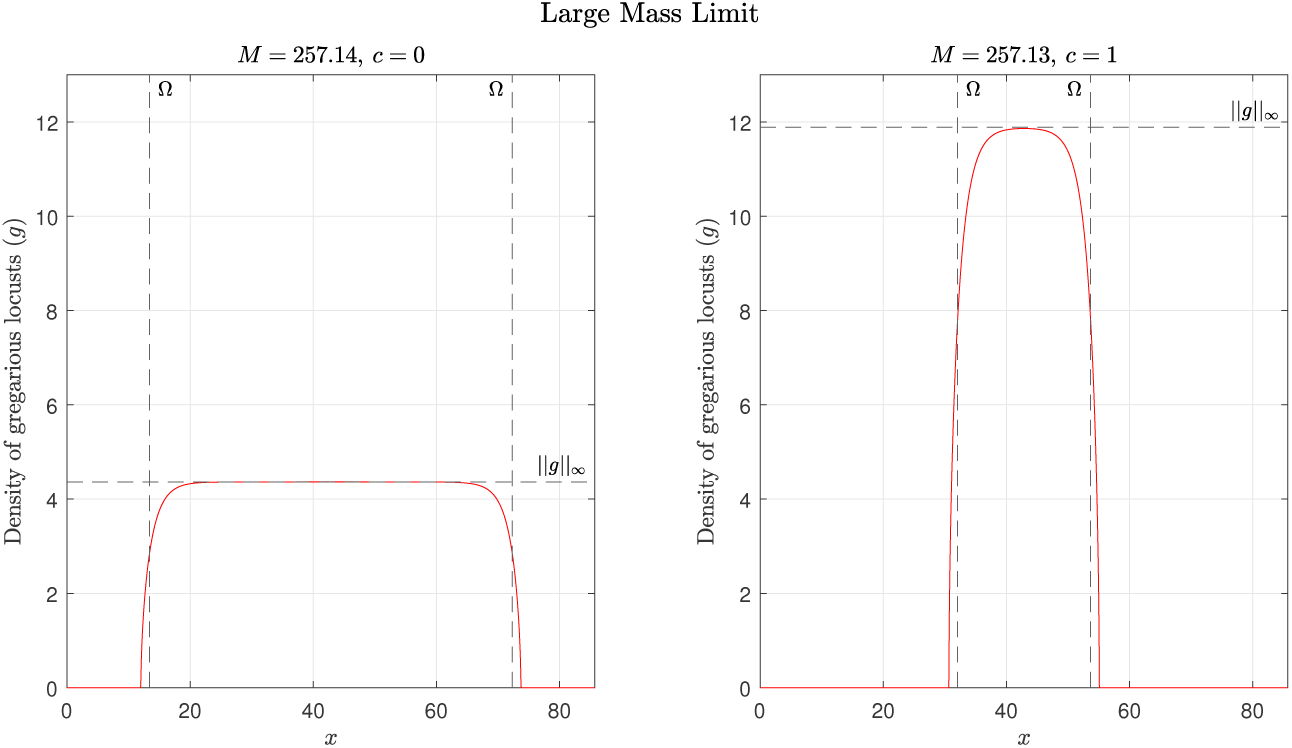
Large mass limit with estimates for the max value and support. The max value and support are labelled ||*g*||_*∞*_ and Ω respectively. For both the simulation and calculations *D* = 0.01, *γ* = 60, *R*_*g*_ = 0.25, *r*_*g*_ = 0.5, *A*_*g*_ = 1, and *c* = 0 and 1. As the mass *M* is increased the gregarious locust shape *g* becomes increasingly rectangular as the maximum locust density does not depend on the total mass. In addition as the amount of food is increased from *c* = 0 on the left to *c* = 1 on the right, the maximum density for the gregarious locusts increases.

For the small mass limit, we begin with Eq (14) and approximate the social interaction potential using a Taylor expansion, 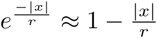 (giving 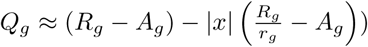. In addition, we ignore the effect of linear diffusion within Ω. These assumptions give Eq (14) as

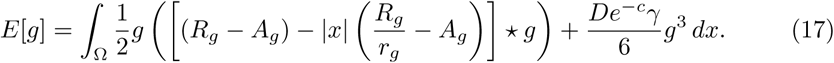

Following [32], gives an estimate of the maximum gregarious locust density, ||*g*||_*∞*_, as

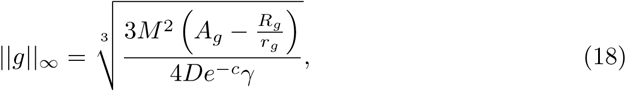

with support,

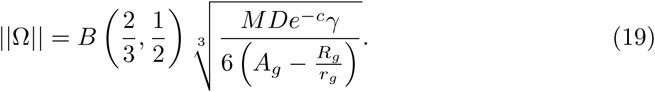

where *B* is the *β*-function (for definition see [36], page 207).

The results of these approximations can be seen in Fig 2. While these approximations are less accurate than those of the large mass limit, they illustrate that as the amount of food increases so too does the maximum hopper band density. However this effect is less pronounced than in the large mass limit. It also demonstrates how the maximum hopper band density and support both increase with an increase in locust mass.

**Fig 2.**
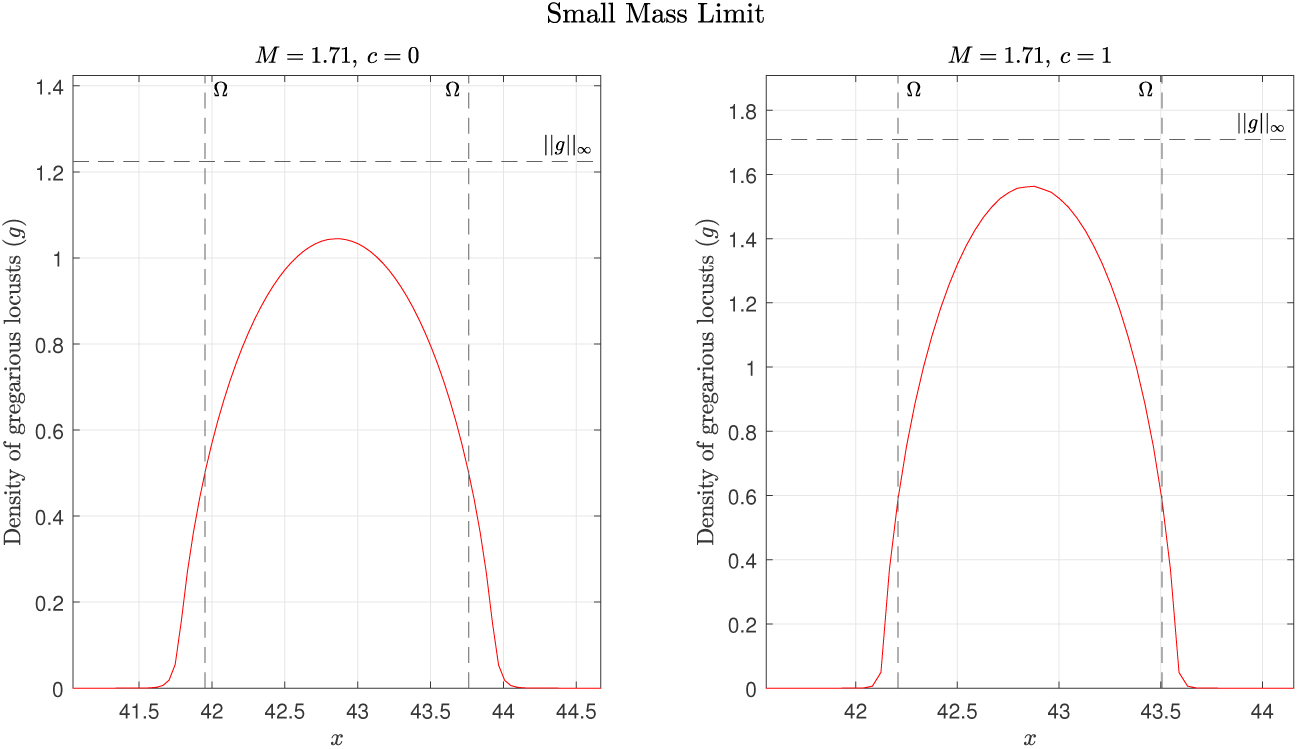
Small mass limit with estimates for the max value and support. The max value and support are labelled ||*g*||_*∞*_ and Ω respectively. For both the simulation and calculations *D* = 0.01, *γ* = 60, *R*_*g*_ = 0.25, *r*_*g*_ = 0.5, *A*_*g*_ = 1, and *c* = 0 and 1.

The accuracy of both the small and large mass approximations and the transition between the two can be seen in Fig 3 for both the maximum hopper band density and support. In the simulations in order to estimate the finite support, Ω, rather than the infinite support due to linear diffusion, *g* = 0.01 was used the cut-off value.

**Fig 3.**
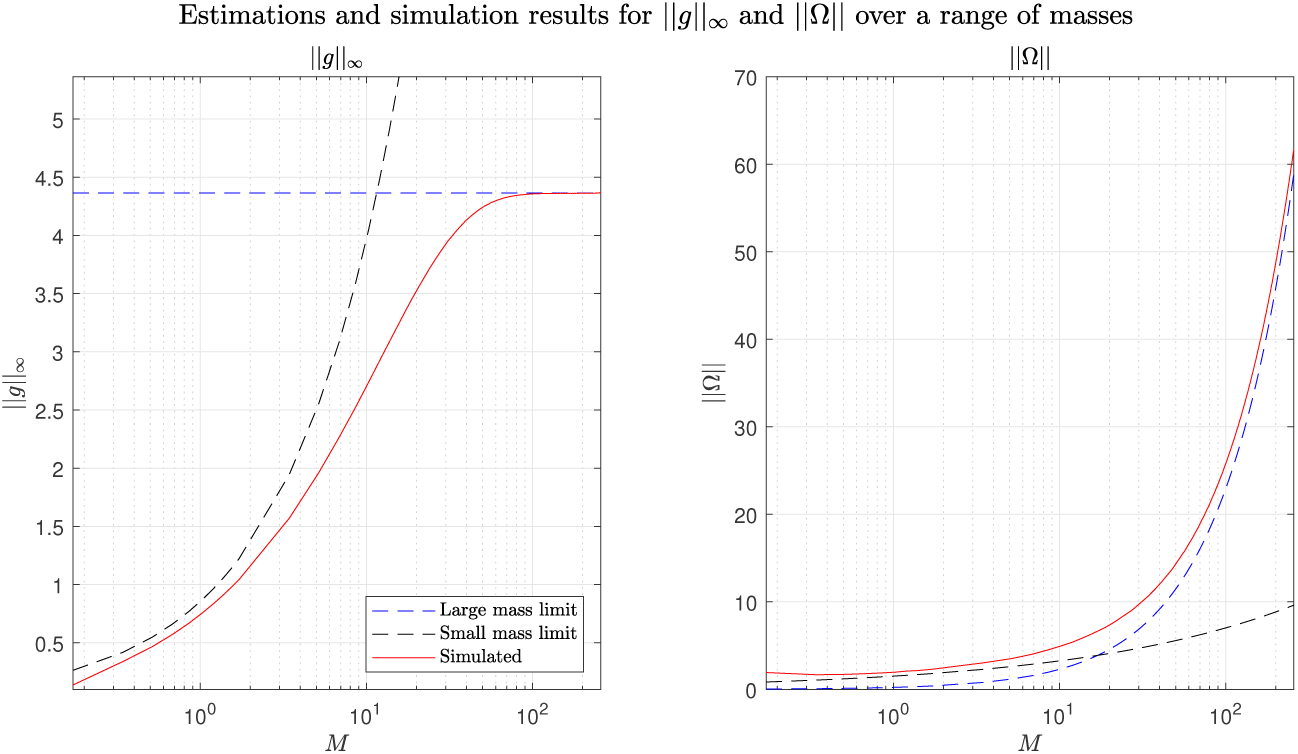
Small and Large mass limit estimates and simulated results for both the maximum hopper band density (left) and support (right). In the simulations in order to estimate the finite support, Ω, rather than the infinite support due to linear diffusion, Ω^*′*^, *g* = 0.01 was used the cut-off value.

#### Linear stability analysis of homogeneous steady states

To see what effect food has on hopper band formation we look at the stability of the homogeneous steady states of *s, g*, and *c*, given by 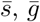, and 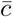, with the total density 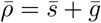. We assume that *c* does not deplete (i.e. *κ* = 0), as otherwise the only homogeneous steady states are 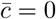 or 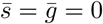. We then perturb the homogeneous steady state to find under what conditions the steady states are unstable. Using this assumption gives the condition for hopper band formation (i.e., instability of the homogeneous steady state) in terms of the global gregarious mass fraction Eq (6) and the total density as

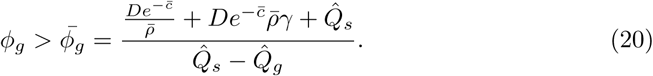

where 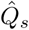 and 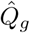 are the Fourier transform of *Q*_*s*_ and *Q*_*g*_ respectively.

From this, it can be seen that as 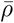 increases the gregarious fraction required for hopper band formation increases suggesting an upper locust density for hopper band formation. However, the effect of 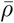 on hopper band formation is diminished as the amount of available food increases.

For our specific functions 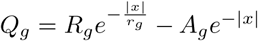 and 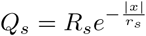, taking the one dimensional Fourier transforms of *Q*_*s*_ and *Q*_*g*_ using the following definition,

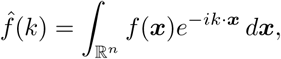

gives the following relationship,

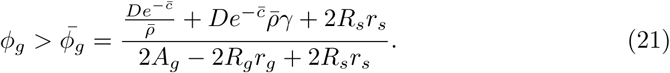

From this we can find the maximum homogenous density, 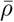, that locust aggregations can still form. So taking Eq (21) and substituting 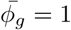 we can solve for 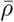 as,

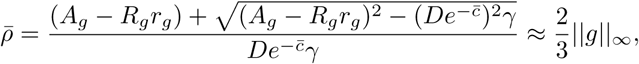

where ||*g*||_*∞*_ is maximum density for the large mass limit given by Eq (15).

Finally, we calculate if it is possible for a particular homogeneous density of locusts to become unstable (and thus form a hopper band). By calculating the homogeneous steady state gregarious mass fraction as,

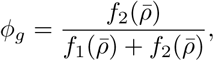

then by combining with (21) we obtain an implicit condition for hopper band formation as

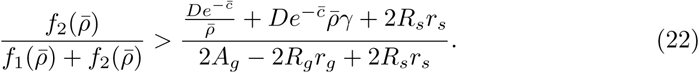

In Eq (22), if the values on the left are not greater than those on the right then it is not possible for a great enough fraction of locusts to become gregarious and for instabilities to occur. As the value of the right hand side decreases as the amount of food increases, we can deduce that the presence of food lowers the required density for hopper band formation.

#### Time until hopper band formation with homogeneous locust densities

We also estimate time until hopper band formation with homogeneous locust densities and a constant *c*. By assuming that *s* and *g* are homogeneous we can ignore the spatial components of Eq (12a) and Eq (12b). We again denote the combined homogeneous locust density as 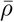 however now 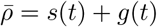. Finally, assuming that *g*(0) = 0, we find the homogeneous density of gregarious locusts as a function of time is given by

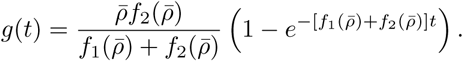

Which we then solve for *t*^*^ such that 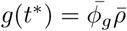, where 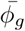 is given by Eq (20). This gives an estimation for time of hopper band formation (i.e. the time required for the homogeneous densities to become unstable) as,

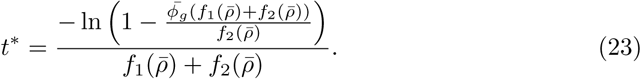

Thus, as increasing food decreases the gregarious mass fraction required for hopper band formation it follows that it also decreases the time required for hopper band formation.

#### Center of mass

Another property of the model is how the center of mass for the locusts behaves. For a single population with diffusive terms it has been shown that the center of mass does not move [29]. Using a similar method we look at how the total population of locusts behaves with a constant food source, i.e. *c*(***x***, *t*) is constant in space and time. We assume that *ρ*(***x***, *t*) → 0 on the boundary of our domain Ω (note: the proof also works if zero flux through the boundary of the domain is imposed), and that *Q*_*s*_ and *Q*_*g*_ are symmetric. For convenience we also let *D*^*^ = *De*^−*c*^. We begin by adding equations (13a) and (13b), and rewrite the result in terms of the local gregarious mass fraction (see Eq (5)), which gives

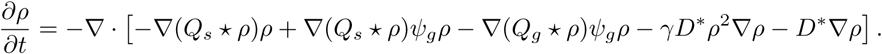

We now consider the behaviour of the center of mass. For notational simplicity we let

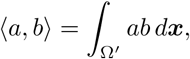

and define the mass of locusts to be

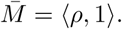

Then the position of the center of mass, *C*, of *ρ*, is given by

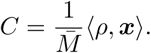

The motion of the center of mass is then given by,

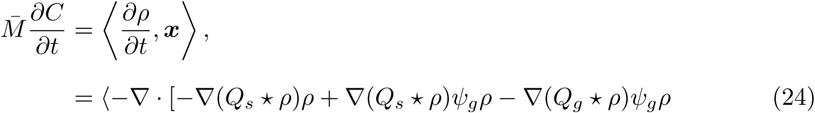

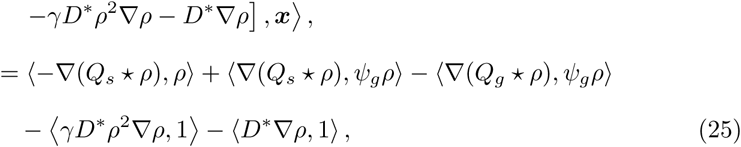

where Eq (25) is obtained using integration by parts and our boundary conditions. Then, considering the diffusion terms in Eq (25), we get

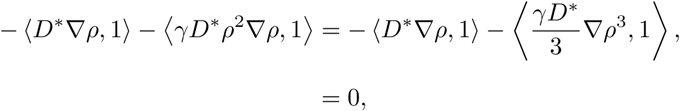

by our boundary conditions and integration by parts. This gives

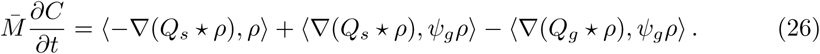

Then using integration by parts and our boundary conditions on Eq (26), we obtain

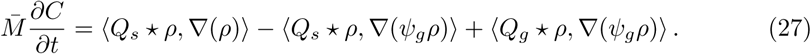

However, using properties of convolutions and the assumption *Q*_*s*_ and *Q*_*g*_ are symmetric, we find and alternate expression for Eq (26) as

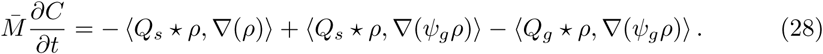

Summing Eq (27) and Eq (28), and dividing by 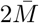 we find

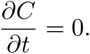

From this we can conclude that in absence of non-uniformities in the food distribution or other movement mechanisms (such as alignment) the center of mass of locusts cannot move.

### Numerical results

We now investigate both the effect of food on locust hopper band formation and the effect of gregarisation on locust foraging efficiency in one dimension. In order to simulate our equations, we used a first order upwinding Finite Volume Scheme for the advection component with Fourier transforms to solve the convolution and central differencing schemes for the diffusion terms. We used an adaptive Runge-Kutta scheme for time, a full detailed derivation can be found in S2 Appendix.

#### Parameter selection and initial conditions

The bulk of the parameters, *R*_*s*_, *r*_*s*_, *R*_*g*_, *r*_*g*_, *A*_*g*_, *k*, and *δ*, have been adapted from [31] to our non-dimensionalised system of equations. We explore two parameter sets that we will term symmetric and asymmetric based on the time frame of gregarisation vs solitarisation. In the symmetric parameter set (*δ* = 1, *k* = 0.681), gregarisation and solitarisation take the same amount of time and the density of locusts for half the maximal transition rate is lower for solitarization. This is the default parameter set from Topaz et. al. [31] with an adjusted *k*_1_ term that is calculated using Eq (22) and the upper range for the onset of collective behaviour as ≈ 65 locusts/*m*^2^ [31, 38]. This behaviour is characteristic of the Desert locust (*Schistocerca gregaria*) [10].

In the asymmetric parameter set (*δ* = 1.778, *k* = 0.1), solitarisation takes an order of magnitude longer than gregarisation, and the density of locusts for half the maximal transition rate is lower for solitarization. This is the alternative set from Topaz et. al. [31]. The Australia plague locust (*Chortoicetes terminifera*) potentially follows this behaviour taking as little as 6 hours to gregarise but up to 72 hours to solitarise [37, 39]. The complete selection of parameters can be seen in Table 1.

**Table 1.** Dimensionless parameters used in numerical simulations for both symmetric and asymmetric gregarisation-solitarisation.

| Variable | Symmetric Value | Asymmetric Value | Source |
| --- | --- | --- | --- |
| $k$ | 0.681 | 0.1 | Eq (22) [10, 23, 31, 37] |
| $\delta$ | 1 | 1.778 | [10, 31, 37] |
| $D$ | 2.041 | 2.041 | |
| $\gamma$ | 431.87 | 294.44 | Eq (15) [17] |
| $R_s$ | 1063.5 | 878.1 | [31] |
| $r_s$ | 1 | 1 | [23, 31] |
| $R_g$ | 940.5 | 775.6 | [31] |
| $r_g$ | 0.2857 | 0.2857 | [23, 31] |
| $A_g$ | 2008.7 | 1658.6 | [31] |
| $\kappa$ | 0.09 | 0.18 | Eq (13c) |

At the densities we are investigating we will assume that the majority of movement will be due to locust-locust interactions rather than random motion, so we set our dimensional linear diffusion term to be of the order 10^−2^, giving our non-dimensional linear diffusion as *D* = 2.041 for both symmetric and asymmetric parametrisation. Next, we estimate the maximum locust density as ≈ 1000 locusts*/m*^2^ [17] and adapt this to our one dimensional simulation as 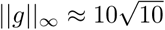 locusts*/m*. Then using Eq [15] we find *γ* = 431.87 for the symmetric parameters and *γ* = 294.44 for the asymmetric parameters.

To estimate *κ* we begin with Eq (13c) and set the nondimensionalised density of locusts to 1 (*ρ* = 1) (and *ρ* = 0.5 for the asymmetric parameters), we then want the locusts to consume approximately 70% of the food over the course of the simulation (i.e., *c* transitions from *c* = 1 to *c* = 0.30)). Solving for *κ* we find *κ* ≈ 0.09 (and *κ* ≈ 0.18 for the asymmetric parameters).

Our spatial domain is the interval *x* = [0, *L*], where *L* = 3*/*0.14, with periodic boundary conditions (i.e., *s*(0, *t*) = *s*(*L, t*)). Our time interval is 12.5 units of time (in dimensional terms this is a 3m domain for a simulated 50 hours).

The initial locusts densities are given by

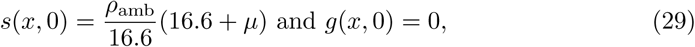

where *ρ*_amb_ is a ambient locust density and *µ* is some normally distributed noise, *µ* ∼ 𝒩 (0, 1). To ensure that simulations were comparable, we set-up three locust initial condition and rescaled them for each given ambient locust density. Finally, the initial condition for food is given by a smoothed step function of the form,

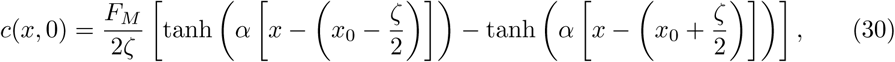

with *α* = 7, *x*_0_ = *L/*2, *F*_*M*_ being the food mass and *ζ* being the initial food footprint. We will also introduce *ω* = 100*ζ/L* which expresses the food footprint as a percentage of the domain.

#### The effect of food on hopper band formation

To investigate the effect that food had on locust hopper band formation, we ran a series of numerical simulations in which the total number of locusts and the size of food footprint were varied, while the total mass of food remained constant. The food footprint ranges from covering 2.5% of the domain to 50% of the domain (*ω* = 2.5 to *ω* = 50). For the symmetric parameters four food masses were tested, *F*_*M*_ = 1.5, 2, 2.5 and 3, and for the asymmetric variables two food masses were tested, *F*_*M*_ = 1.5 and 3. As a control we also performed simulations with both no food present and a homogeneous food source, represented by *ω* = 0 and *ω* = 100 respectively, for each ambient locust density.

We varied the ambient locust density ranging from *ρ*_amb_ = 0.8 to *ρ*_amb_ = 1.4 for the symmetric parameters. This range was selected based on Eq (22) so that in the absence of food hopper band formation would not occur. We also ran three simulations for each combination of *ρ*_amb_, *ω,* and *F*_*M*_ with varied initial noise and took the maximum peak density across the three simulations, as we found in certain cases the initial noise had an effect on whether a hopper band would form.

For the asymmetric variables we varied *ρ*_amb_ from *ρ*_amb_ = 0.3 to *ρ*_amb_ = 0.55, to test the effect food had on the time frame of hopper band formation. From Eq (22) there should be hopper band formation at the upper half of this density range, however from Eq (23) this will only occur outside or right at the end of our simulated time frame. We ran a single simulations for each combination of *ρ*_amb_, *ω,* and *F*_*M*_.

The results for the symmetric parameter experiments are displayed in Fig 4. The plots show the final peak gregarious density for the varying food footprint sizes and ambient locust densities. In the blue regions there was no hopper band formation and in the green regions there was successful hopper band formation. It can be seen in the plots that as the food mass is increased the minimum required locust density for hopper band formation decreases. This effect is more pronounced within an optimal food width and this optimal width increases as as the amount of food increases.

**Fig 4.**
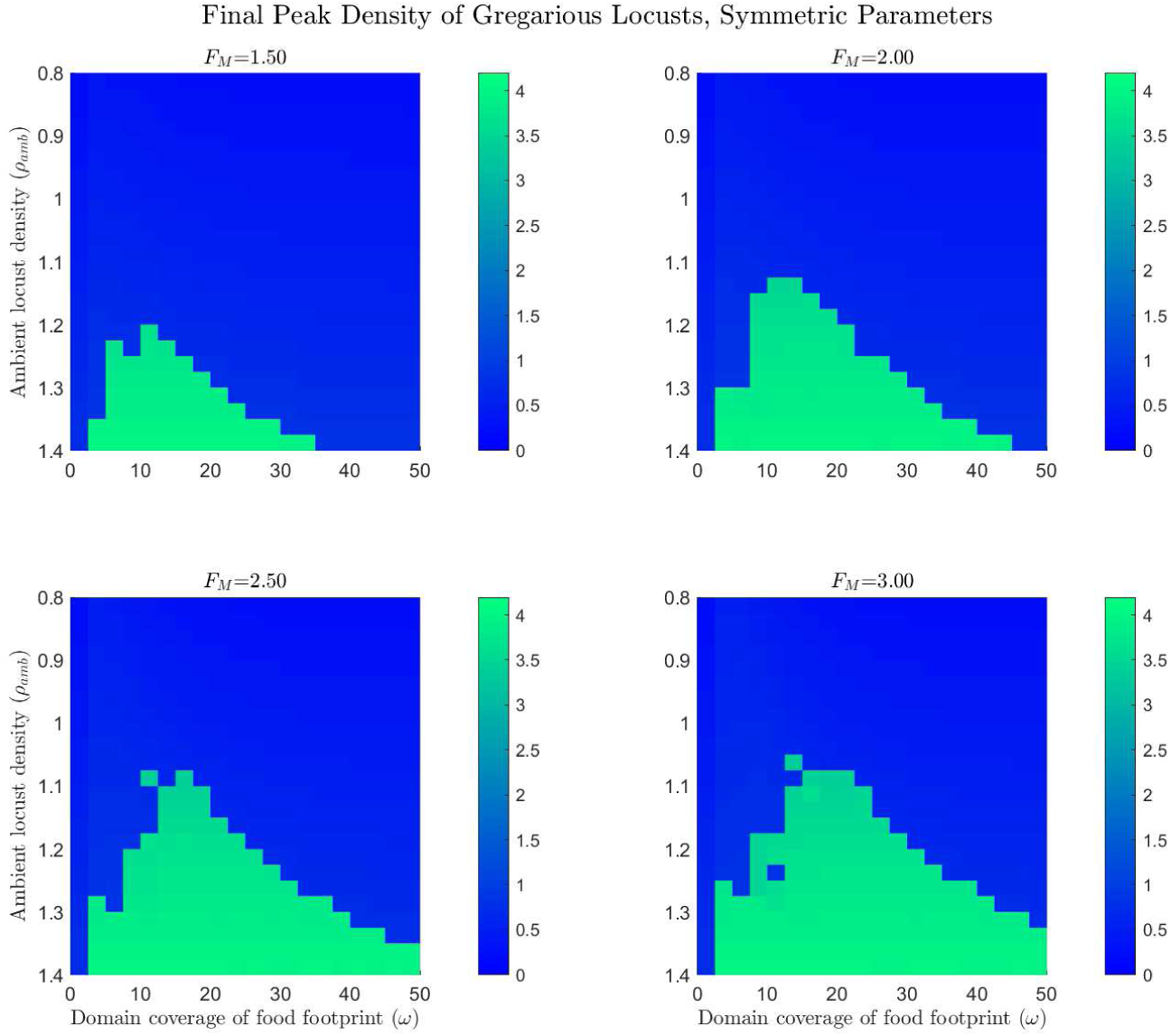
Maximum gregarious locust density for the symmetric gregarisation parameters with varying food footprint sizes and initial ambient locust densities. For the simulations, *x* = [0, 3*/*0.14] with periodic boundary conditions and *t* = [0, 12.5]. The initial condition for locust densities is given by Eq (29) and food initial conditions are given by Eq (30). Ambient locust density ranges from *ρ*_amb_ = 0.8 to *ρ*_amb_ = 1.4, food footprint ranges from *ω* = 0 to *ω* = 50, the food mass *F*_*M*_ = 1.5, 2, 2.5 and 3, and the consumption rate *κ* = 0.09. The plots show the final peak gregarious density for the varying food footprint sizes and ambient locust densities, in the blue regions there was no hopper band formation and in the green regions there was successful hopper band formation.

The results for the asymmetric parameter experiments are displayed in Fig 5. Again, green indicates successful hopper band formation and blue indicates no hopper band formation. It can be seen in these plots that with no food present a hopper band failed to form within the simulated time. From this we can infer that food also decreases the required time for hopper band formation, again there is an optimal food width for this effect.

**Fig 5.**
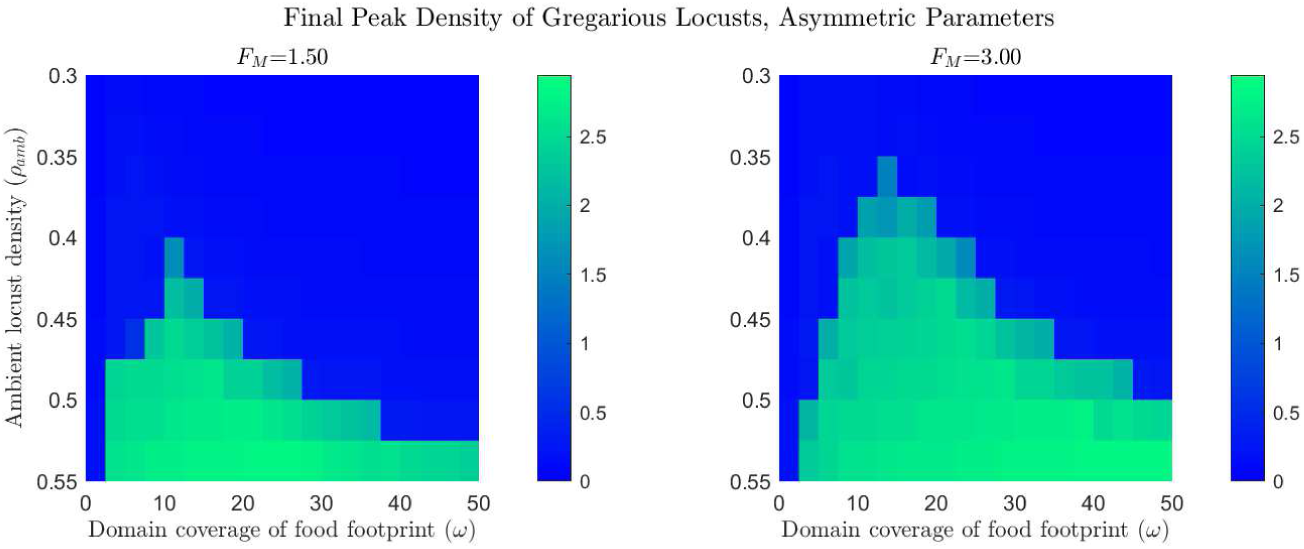
Maximum gregarious locust density for the asymmetric gregarisation parameters with varying food footprint sizes and initial ambient locust densities. For the simulations, *x* = [0, 3*/*0.14] with periodic boundary conditions and *t* = [0, 12.5]. The initial condition for locust densities is given by Eq (29) and food initial conditions are given by Eq (30). Ambient locust density ranges from *ρ*_amb_ = 0.3 to *ρ*_amb_ = 0.55, food footprint ranges from *ω* = 0 to *ω* = 50, the food mass *F*_*M*_ = 1.5 and 3, and the consumption rate *κ* = 0.18. The plots show the final peak gregarious density for the varying food footprint sizes and ambient locust densities, in the blue regions there was no hopper band formation and in the green regions there was successful hopper band formation.

We can delve deeper into the results by looking at a representative sample of simulations in Fig 6. In these simulations *ρ*_*amb*_ = 1.2, *κ* = 0.09, and *F*_*M*_ = 1.5, with food footprints *ω* = 7.5, 10, and 12.5 as well as with no food present. In the simulations in which food is present, gregarious locusts aggregate at the center of the food. If the food source is too narrow (*ω* = 7.5, *t* = 3) there is an attempt at hopper band formation but the gregarious mass is too small and the food source has not been sufficiently depleted so a large portion remains within the food source, thus the hopper band does not persist. If the food is too wide (*ω* = 12.5) the gregarious locusts simply cluster in the center of the food and don’t attempt hopper band formation. However, if the food width is optimal (*ω* = 10) there is a successful hopper band formed, this is seen as clump or aggregation of gregarious locusts in the final plot.

**Fig 6.**
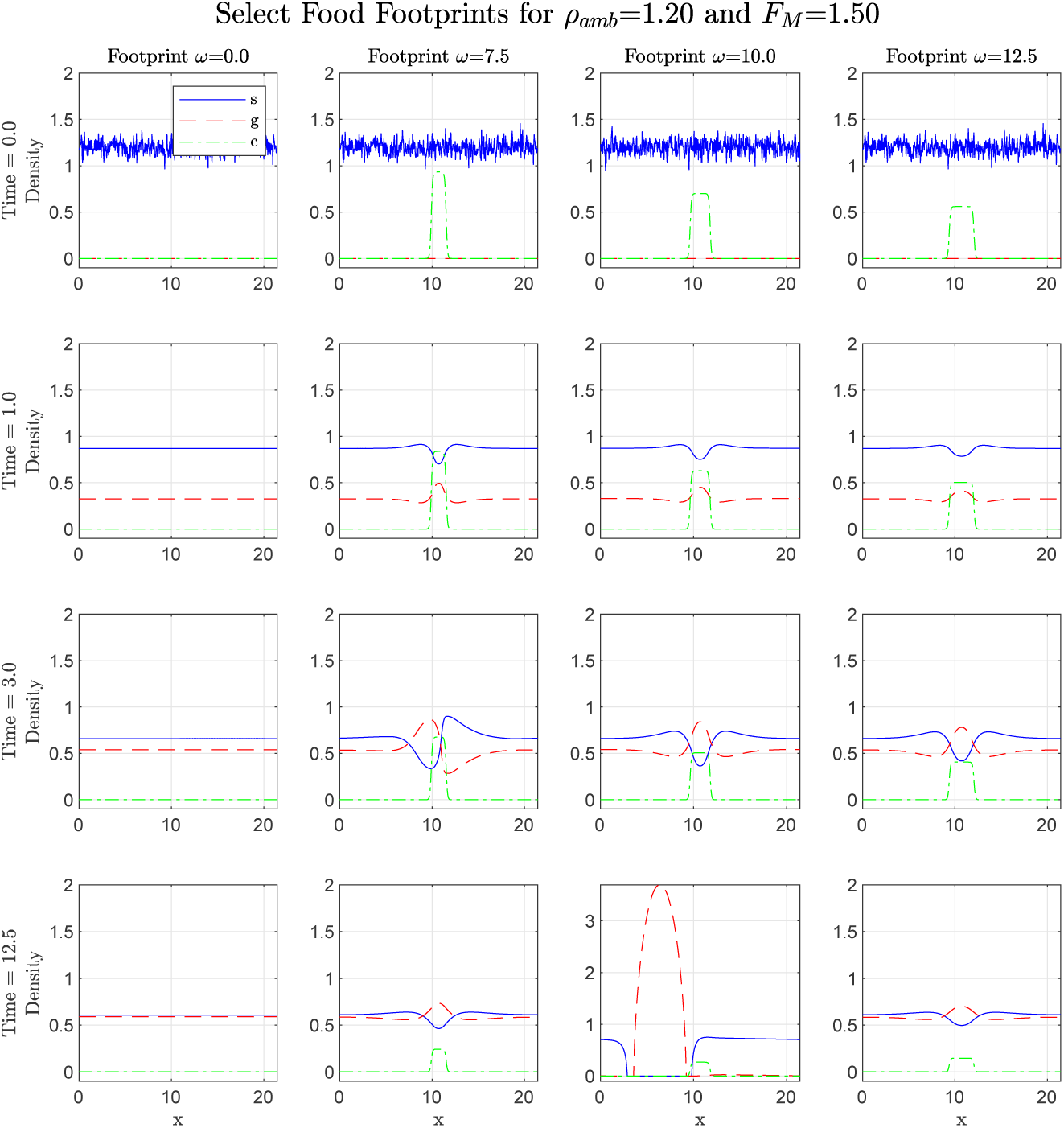
A selection of plots showing the effect of food distribution on gregarisation and locust hopper band formation. In these simulations *ρ*_*amb*_ = 1.2, *κ* = 0.09, and *F*_*M*_ = 1.5 with *ω* = 7.5, 10, and 12.5, as well as with no food present (labelled *ω* = 0).

#### The effect of gregarisation on foraging efficiency

It is also possible to investigate the effect of gregarisation on foraging efficiency. Using [40] as a guide we first calculate the per capita contact with food for solitarious and gregarious locusts, respectively as

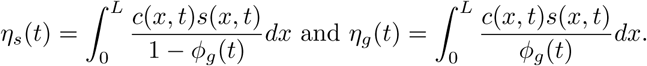

We then calculate the instantaneous relative advantage at time *t* as

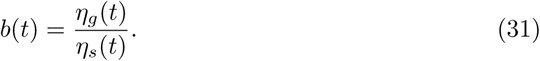

By looking at the instantaneous relative advantage vs the global gregarious mass fraction prior to hopper band formation in Fig 7, it can be seen that as the gregarious mass fraction increases so too does the foraging advantage of being gregarious. Thus, as a greater proportion of locusts become gregarized it is more advantageous to be gregarious. This effect is increased by the mass of food present but is diminished by the size of the food footprint to the point where no advantage is conferred when the food source is homogeneous. This effect is visualised in Fig 6, as gregarious locusts aggregate in the center of the food mass and displace their solitarious counterparts.

**Fig 7.**
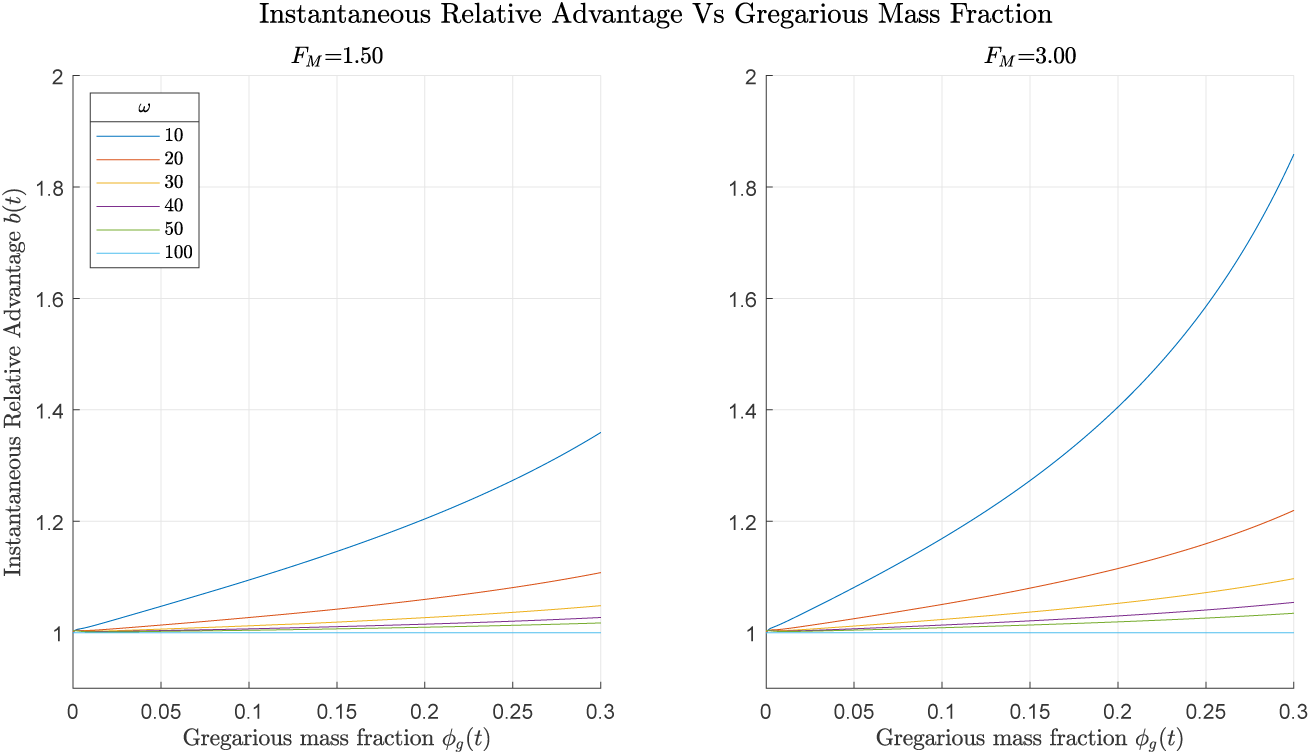
Instantaneous relative advantage of gregarious locusts vs gregarious mass fraction at various food footprints and food masses. In these simulations *ρ*_*amb*_ = 0.95 and *κ* = 0.09. The homogeneous food source is labelled *ω* = 100. It can be seen that as the gregarious mass fraction increases so too does the foraging advantage of being gregarious, this effect is increased by the mass of food present but is diminished by the size of the food footprint.

## Discussion

Locusts continue to be a global threat to agriculture and food security, and so being able to properly predict and control outbreaks is of great importance. In this paper we presented a continuum model that includes non-local and local inter-individual and interactions with food resources. This model extends the existing Topaz et. al. 2012 [31] model for locust gregarisation to include food interactions and local repulsion. By analysing and simulating our new model we have found that food acts to: increase maximum locust density, lower the gregarious fraction required for hopper band formation, and decreases both the required density and time for hopper band formation with this effect being more pronounced at some optimal food width.

Analytical investigations of our model shows that a spatially uniform food source has a variety of effects has on locust behaviour. Firstly, by considering a purely gregarious population we found that the maximum locust density is affected by the amount of food present, in that increasing food leads to increased maximum density. Then, by performing a linear stability analysis we found the gregarious mass fraction required for hopper band formation depends on both the ambient locust density and the amount of food present, with increasing food decreasing the required gregarious mass fraction. Using this relationship we then found that our model also has a theoretical maximum locust density for hopper band formation, and that the presence of food lowers both the required time and density of locusts for hopper band formation. Finally, we have also shown that the center of mass of locusts is not dependent on the locust-locust interactions we explored, so prior to extra interactions such as alignment locusts will aggregate at roughly the center of food sources. This leads to a possible extension of the model by including the alignment component of locusts collective movement.

While it has been shown that highly clumped food sources lead to a greater likelihood of gregarisation [20], using numerical simulations we have shown that there exists an optimal width for these food clumps for hopper band formation. This effect was shown to lower the required density for hopper band formation via the symmetric parameters and the required time via the asymmetric parameters. This optimal width is dependent on the amount of food present relative to the locust population. This effect appears to be brought about by the depletion of the food source, if the food source is not sufficiently depleted, then a gregarious hopper band will fail to form because a portion of the gregarious population will remain on the food.

In 1957 Ellis and Ashall [41] found that dense but patchy vegetation promoted the aggregation of hoppers and that sparse uniform plant cover promoted their dispersal. By looking at the relative foraging advantage of gregarious locusts in these situations we found that as the gregarious mass fraction increases so too does the foraging advantage of being gregarious. This effect is increased by the mass of food present but is diminished by the size of the food footprint to the point where no advantage is offered with a homogenous food source. While there are various explanations about the costs and benefits of group living [42], there are very few explanations about the evolution of phase polyphenism (but see the predator percolation hypothesis in [43]) and why some animals would switch back and forth between solitary and gregarious phenotypes. Our results, in line with recent studies about solitary and social foraging in complex environments [44], provide a possible evolutionary explanation for Ellis and Ashall observations.

Preventative methods are the key to improving locust control. This includes the ability to predict mass gregarisation according to resource distribution patterns so that the area searched for locusts is reduced and control efforts are deployed in high risk areas early on [18]. Our results have the potential to improve predictive gregarisation models and early detection efforts by further increasing our understanding of the link between gregarisation and vegetation (resource) distribution, the latter becoming increasingly easy to quantify during field surveys, and aerial surveys including drones and satellite imagery [21, 37]. Future research should focus on developing decision support systems integrating predictive gregarisation models and GIS data from surveys.

## Acknowledgements

The authors would like to thank Andrew J. Bernoff for the insightful discussions.

## Supporting information

**S1 Appendix Detailed analytic results.** The full detailed derivations of the analytic results given in the PDE model analysis section.

**S2 Appendix Numerical Scheme.** The full detailed derivation of the numerical scheme used for simulating the numerical results.

